# Theta and gamma transcranial alternating current stimulation modulate Mandarin consonant and lexical tone perception

**DOI:** 10.1101/2025.09.30.679546

**Authors:** Yaxuan Wang, Keke Yu, Shuqi Yin, Baishen Liang, Ruiming Wang

## Abstract

A theta/gamma oscillatory neural mechanism has been postulated to explain the auditory sampling of hierarchical syllable-phoneme structure with corresponding speech rates or linguistic hierarchies (theta for syllable and gamma for phoneme). Yet, whether such a mechanism is generalizable to Mandarin Chinese, a language that both has suprasegmental phonemes in the form of lexical tones and contains more monosyllabic words, which may lead to higher reliance on either phonemic-level or syllabic-level perceptual representations, is unclear. In this study, we applied transcranial electric stimulation with either theta or gamma alternating currents (tACS) to bilateral auditory cortices in healthy Mandarin-speaking participants during a consonant or tone identification task in quiet or noisy environments. Results showed that theta tACS impaired consonant identification in quiet, specifically, and generally prolonged reaction times across tasks and environments. Gamma tACS, however, only delayed tone identification in quiet. Besides, both theta and gamma tACS modulated perceptual decision-making parameters, leading to increased boundary thresholds (cautiousness of decision) and altered response biases in the perceptual decision of both consonants and tones, as evidenced by the hierarchical drift-diffusion model (HDDM). These findings indicate that theta oscillations causally support the perception of Mandarin syllables, with consonants and lexical tones presumably embedded in syllable-level representations, regardless of task difficulty. Gamma activity, however, is presumably engaged in supporting fast-changing and fine-grained acoustic features in a modulatory manner.

**Significance Statement:** How the brain samples speech features for perception is a key question in language neuroscience. Previous research has shown that theta and gamma brain activity support the perception of suprasegmental syllables and segmental phonemes, corresponding to their length or linguistic hierarchy. However, for Mandarin, the existence of suprasegmental phonemes (lexical tones) and the large number of monosyllabic words can either lead to higher reliance on gamma oscillations for phoneme perception, or higher dependence on theta oscillations for holistic syllable integration. With the combination of brain stimulation to selectively modulate theta/gamma auditory activity and phoneme identification tasks, this study reveals a causal role of theta activity in supporting holistic syllable-based Mandarin phoneme perception, yet gamma engagement is modulatory.

## Introduction

Spoken Mandarin relies on the perception of syllables, which integrate phonemic features such as consonants and lexical tones (Liang & Du, 2018). This requires the auditory neural system to sample speech at multiple timescales. Neural oscillations are key in tracking and sampling speech signals (Giraud & Poeppel, 2012; Meyer, 2018). Especially, neural oscillations in the theta and gamma bands are shown to support speech sampling by aligning with distinct acoustic structures.

Theta oscillations (4-8Hz) have been found to sample slow-changing suprasegmental cues (Meyer, 2018), including syllables, phrases, and prosodic cues (e.g., intonation) (Teoh et al., 2019), as well as lexical tones (Wang et al., 2017). In Mandarin, which has more monosyllabic semantic units than English (Kessler & Treiman, 1997; Wu et al., 2023), theta oscillations also play a key role by adaptively tracking syllabic speech rhythms to optimize perception (He et al., 2023; Ni et al., 2023). Evidence from event-related potential (ERP) (Zou et al., 2020) and articulatory phonetics (Kang & Xu, 2024; Liu & Xu, 2023) studies further suggests that lexical tones and segmental phonemes are processed in temporal synchrony, indicating that syllable perception in Mandarin depends on the coordinated timing of their phonemic components.

Complementing this, gamma oscillations are involved in processing fine-grained phonemic features varying rapidly in time, including formant transitions (Fukuda et al., 2010) and voice onset times (Rufener, Oechslin, et al., 2016), which are crucial for differentiating meanings (e.g., bear vs. pear). Besides, recent transcranial alternating current stimulation (tACS) studies modulating neural oscillations (Ghiani et al., 2021; Herrmann et al., 2016) have shown that theta and gamma stimulation selectively perturb neural tracking of syllable rhythms (Zoefel et al., 2018) and consonant discrimination (Rufener, Oechslin, et al., 2016), respectively. In brief, theta oscillations support global syllabic integration, whereas gamma oscillations resolve local phonetic features (Morillon et al., 2012; Lizarazu et al., 2019).

In addition, such a global-local theta/gamma sampling mechanism can be modulated in adverse listening conditions (Kegler & Reichenbach, 2021; Keshavarzi et al., 2020). Electroencephalogram (EEG) studies showed that theta activity is enhanced in speech-in-noise tasks, whereas gamma activity is maintained under clear speech conditions for fine-grained structure analysis (Yellamsetty & Bidelman, 2018).

Nevertheless, as most studies focused on non-tonal and multisyllabic languages, it remains unclear whether and how theta/gamma sampling applies to tonal and relatively monosyllabic languages like Mandarin (Liang & Du, 2018). Specifically, whether lexical tones are encoded as segmental phonemes (gamma-based), as both determine meanings, or suprasegmental prosodies (theta-based), as both fluctuate in syllable length, is unclear. Moreover, it remains elusive whether Mandarin’s monosyllabic nature promotes holistic syllable-level encoding or retains the nested theta/gamma structure.

Here, we investigate the causal roles of theta/gamma oscillations in Mandarin syllable perception and their modulation by noise through an online tACS study combined with a Mandarin phoneme identification task. Theta or gamma tACS, found able to perturb the corresponding bands of neural oscillations (de Lara et al., 2018; Ghiani et al., 2021), was applied to bilateral temporal areas while participants were identifying the consonant or lexical tone of resynthesized, perceptually ambiguous syllables drawn from 5-step continua (i.e., [t]-[t^h^] and tone1-tone2). This paradigm helps detect stimulation-induced changes in the perceptual sensitivity of speech categories (D’Ausilio et al., 2009; Liang et al., 2023), revealing whether theta/gamma oscillations support Mandarin syllable perception in a frequency-specific manner.

We propose three competitive hypotheses regarding how theta/gamma oscillations might support Mandarin syllable perception (Fig. 1). Hypothesis 1 posits that theta and gamma oscillations sample suprasegmental (e.g., tones) and segmental (e.g., consonants) features, respectively, based on their acoustic length or linguistic hierarchy. Hypothesis 2 suggests that both consonants and tones are encoded as phonemes, with gamma oscillations playing a dominant role. Hypothesis 3 proposes that lexical tones and consonants are encoded as integrated syllabic units in perception, subserved by theta oscillations. Additionally, considering adverse listening conditions, based on previous findings, we also hypothesize that theta oscillations become more essential in speech-in-noise tasks.

**Figure 1.**
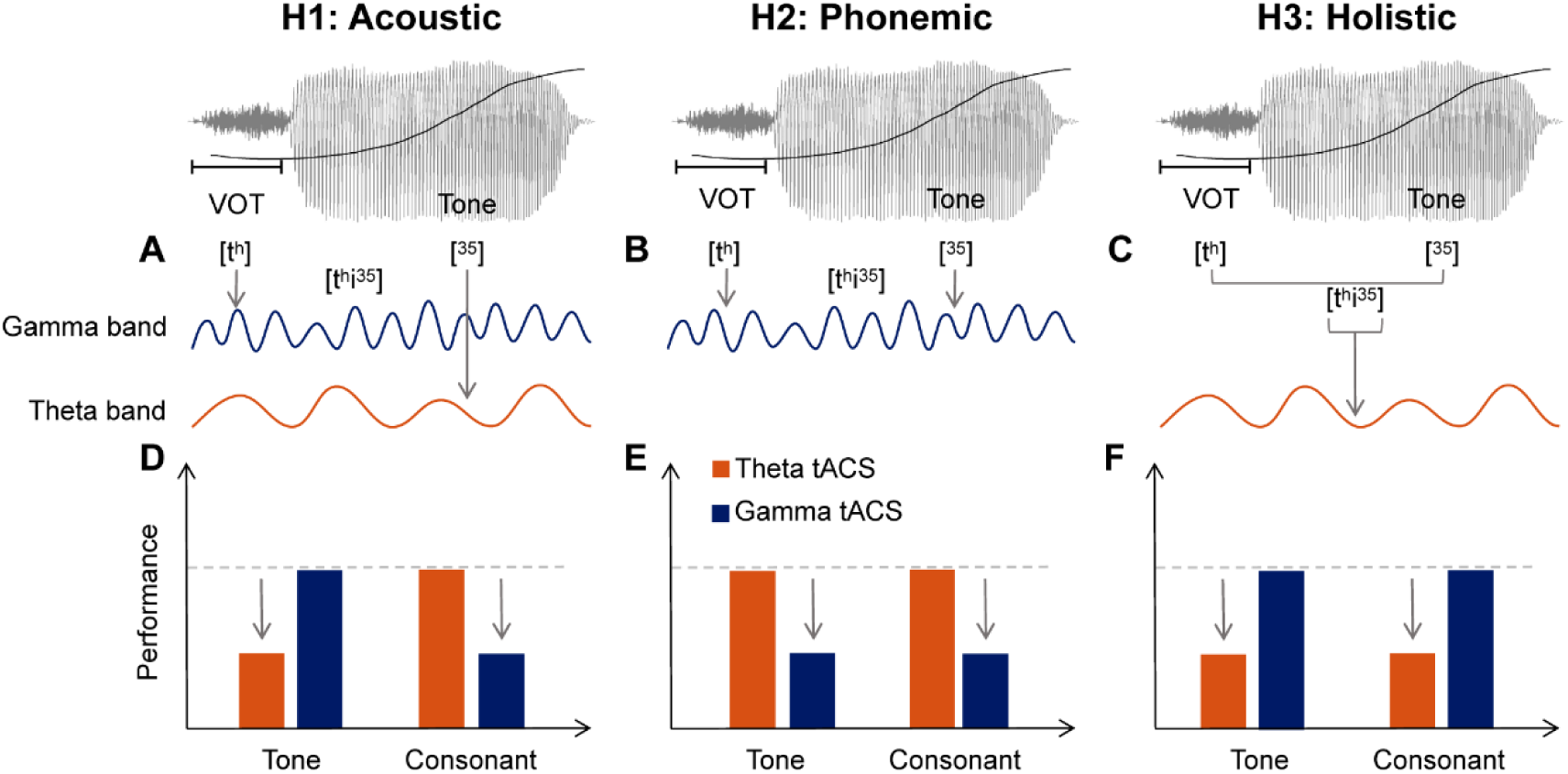
Three proposed conceptual models of Mandarin syllable perception and the predicted effects of tACS. The acoustic structure of a sample Mandarin syllable (e.g., “tí”, [t^h^i^35^]), showing the voice onset time (VOT) and the fundamental frequency (F0) contour. ***A-C,*** Schematic illustrations of the gamma and theta oscillatory neural mechanisms proposed by each hypothesis to process the syllable. ***D-F,*** Predicted patterns of performance impairment on consonant and tone identification tasks under theta and gamma tACS, relative to a sham stimulation baseline (gray dotted horizontal lines). Hypothesis 1 (the Acoustic Hypothesis) proposes a distinctive role of gamma and theta oscillations to support consonant and lexical tone perception, respectively (***A***), which predicts a selective impairment by gamma tACS on consonant perception, and theta tACS on tone (***D***). Hypothesis 2 (the Phonemic Hypothesis) posits that both consonants and tones are represented as phonemic units primarily supported by gamma oscillations at phoneme rates (***B***), which predicts a gamma-specific disruption: gamma tACS should impair performance in both consonant and tone perception, whereas theta tACS is expected to exert little influence (***E***). Hypothesis 3 (the Holistic Hypothesis) proposed that Mandarin consonants and lexical tones are embedded in integrated syllabic representations during perception, with theta oscillations at syllable rates serving as the main neural mechanism (***C***). This predicts a general deficit under theta tACS across both consonant and tone tasks, while gamma tACS is not expected to produce marked effects (***F***).

## Materials and Methods

### Participants

We initially recruited 35 adult native Mandarin speakers aged 18-25 years (7 male, mean age = 20.57 years, SD = 1.93), all of whom were undergraduate or graduate students in Guangzhou with higher education. 30 of these participants (6 male, mean age = 20.67 years, SD = 1.84) who fulfilled the recruiting criteria were included in the final analysis. The group size was sufficient to detect medium to significant effects, as estimated by G*Power 3.1.

All participants were right-handed, as assessed by a modified version of the Edinburgh Handedness Inventory (mean laterality quotient = 101.09, SD = 20.86). In this modified version, the original response options were simplified into three categories: left hand (−2 points), both hands (0 points), and right hand (2 points), with scoring methods consistent with the original inventory (see Supplementary Materials, Q3). Participants all had normal hearing, as measured using customized MATLAB scripts with a pair of Edifier MR4 Monitor Speakers at frequencies of 0.25, 0.5, 1, 2, 3, 4, 6, and 8 kHz (≤ 25dB SPL) in a soundproof room. None of them had a self or family history of neurological, traumatic, or psychiatric disease. 4 of the 30 participants had received formal musical training that had lasted longer than two years. All participants gave written informed consent before the experiment and were paid 40 yuan per hour. The experimental protocol (SCNU-PSY-2024-266) was approved by the Human Research Ethics Committee for Non-Clinical Faculties of the School of Psychology, South China Normal University.

Five participants were excluded from the final sample for the following reasons: difficulties during the task of consonant perception (n = 1), or ineligibility (n = 4) due to current neurological or psychological disorders, a history of seizures, or a score higher than 25 dB from 0.25 to 8 kHz.

### Acoustic Stimuli

Syllables used in this experiment are four natural Mandarin monosyllables: [ti^55^], [ti^35^], [t^h^i^55^], and [t^h^i^35^], transcribed using both the International Phonetic Alphabet and Chao’s “tone-letters” system (Liang et al., 2023). These corresponded to Mandarin syllables “堤dī” (riverbank), “敌dí” (enemy), “梯tī” (ladder), and “题tí” (title), respectively, in Pinyin. Syllables were low-pass filtered at 5 kHz using Praat 6 algorithms and matched for average root-mean-square (RMS) sound pressure level (SPL). To account for acoustic features, the four syllables were denoised, re-synthesized, and pitch contours normalized using Praat 6. In the pretest of the experiment, individualized tone continua ([ti^55^] to [ti^35^]) and consonant continua ([ti^55^] to [t^h^i^55^]) were generated based on four natural syllables by stepwise adjustment of F0 and addition of aspiration noise, with each continuum consisting of five steps (Fig. 2*G*). This approach ensured a range of categorical clarity across the stimuli, as categorically quiet speech has been shown to exhibit greater resistance to noise interference compared to ambiguous speech categories (Bidelman et al., 2020). Crucially, the phonetic continua allow for a comprehensive examination of perception under varying noise conditions. Midpoints in the continuum, rather than eliciting ceiling or floor effects, provide an optimal window to detect neuromodulation effects, as small perceptual biases induced by tACS can be more readily observed. This design thus enables a more precise assessment of neuro-modulatory effects across both quiet and noisy contexts.

**Figure 2.**
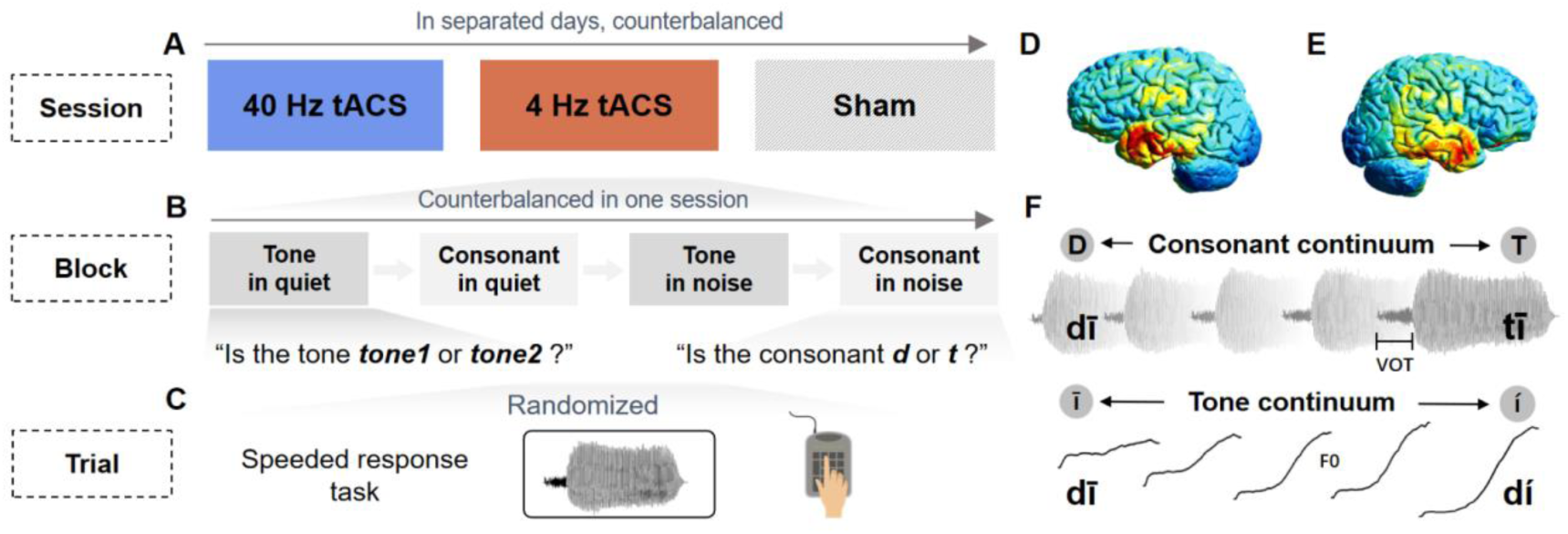
Experiment design, tACS montage, and stimulus continua. ***A-C****, Design of the experiment. **A***, Participants received different patterns of tACS in different sessions with order counterbalanced: gamma band (40 Hz) tACS, theta band (4 Hz) tACS, and sham. ***B,*** One session included four blocks with the order counterbalanced: lexical tone and phoneme (consonant) identification tasks. ***C***, Participants made speeded decisions. The experimental stimuli consisted of four syllables created by crossing two tones (tone 1 [^55^] and tone 2 [^35^] with two consonants ([t] and [t^h^]), resulting in four unique syllables: [ti^55^](堤, riverbank), [ti^35^] (敌, enemy), [t^h^i^55^](梯, ladder), and [t^h^i^35^](题, title). ***D-E****, TACS montage.* Theta or gamma band tACS was applied to bilateral temporal areas by sponge electrodes placed on T7 and T8 (10-20 EEG montage). ***D***, Left view. ***E***, Right view. Voltage maps were simulated by the SimNIBS software (https://simnibs.github.io/simnibs/build/html/tutorial/gui.html). Note that the polarities and values alternated over time. ***F***, *Phonetic continua.* Two types of five-step continua were synthesized. The consonant continuum (top) varied in voice onset time (VOT) from [t] to [t^h^]. The tone continuum (bottom) varied in fundamental frequency (F0) from Mandarin tone 1 (high level) to tone 2 (rising).

The auditory stimuli were delivered using Edifier MR4 Monitor Speakers in stereo. A Deli DL333202 decibel meter (A-weighted) was employed to measure and calibrate the intensity levels of stimuli. In this experiment, each syllable was presented at a SPL of 65 dB through Psychtoolbox-3 in MATLAB R2016a.

In noise conditions, speech-spectrum noise (SSN) was played in synchrony with syllables. SSN was generated by spectrally modulating white noise using the long-term average spectrum of 5000 Mandarin sentences produced by 10 young female speakers, followed by low-pass filtering at 5 kHz (Wu et al., 2005). For each participant, the intensity of SSN was adjusted individually to produce signal-to-noise ratios (SNRs) at a moderate level in a pretest before the experimental session.

### TACS Stimuli

Two rubber electrodes (35 cm^2^) covered by sponge pads and wetted with a 0.9% saline solution (5 mL per electrode) were placed on the T7 and T8 locations (International 10-20 system of electrode placement, Fig.2*D-F*). The electrodes were connected to a neurostimulation device (Soterix Medical 1 ×1 Transcranial Electrical Stimulation Model 2001 Low-Intensity Stimulator). The resistance between the electrodes of each device was set to be no more than 10 kΩ. For each electrode, the stimulation intensity was set at an individualized level such that participants felt comfortable and uncertain about the presence of electric stimulation.

To determine the maximum magnitude of current stimulation for each participant, two sets of 5s sinusoidal testing currents with frequencies of 4 Hz and 40 Hz were applied, respectively. The signal amplitude was measured in a staircase manner, starting at 0.8 mA and adjusted in steps of 0.1 mA up to a maximum of 1.5 mA. The amplitude adjustment was stopped once the subject reported experiencing a skin sensation, and the final amplitude was recorded as the maximum magnitude of current stimulation for subsequent study sessions. Across participants, we therefore obtained a mean amplitude of 0.83 mA for theta tACS (SD = 0.14 mA) and 1.02 mA for gamma tACS (SD = 0.17 mA).

### Experimental procedure

All participants undertook four experimental sessions on four separate days. Each participant came four times to finish one pretest session and three formal experimental sessions, respectively. Participants were seated in a soundproof room. The pretest began with questionnaire surveys (see Supplementary Materials Q1-4), including an informed consent form, demographic surveys, music experience evaluations, and a handedness inventory. Following this, participants completed a pure tone average test and an electric stimulation toleration test. Subsequent psychoacoustic tests first established individualized ambiguity ranges for pitch and voice-onset time (VOT) continua in quiet conditions, followed by a syllable-in-noise perception task to determine individualized perceptual thresholds for these features in noisy environments.

To estimate the individualized perceptual ambiguity ranges for tone and consonant continua, participants undertook tone or consonant identification tasks using a 9×9-step subset from a 54×54-step continuum matrix, applying the method of constant stimuli. During each trial, participants listened to a syllable and were required to identify the tone or consonant category to which it belonged, disregarding the other dimension. The boundaries of the perceptual ranges were manually identified by experimenters. Based on the individualized parameters, we synthesized one 5-step lexical tone and one 5-step consonant continuum for each participant. Re-synthesized syllables were longer than a period of theta oscillation (mean = 0.378s > 0.250s, SD = 0.005s), but a theta window covers enough time samples for consonant or lexical tone recognition, so we would not scale the syllables to sacrifice authenticity. Nevertheless, differences in VOT length between step 1 and step 3 syllables surrounding the point of subjective equity (PSE) were similar to a gamma period (mean = 0.020s ≈ 0.025s, SD = 0.005s), matching the target band modulated by gamma tACS.

Individualized SNR levels were also estimated via the constant stimuli method, where participants performed tone and consonant identification tasks utilizing clear syllables ([ti^55^] and [ti^35^] for tones, [ti^55^] and [t^h^i^55^] for consonants), obscured by SSN. The SSN was presented in synchronization with the syllables with 10-ms linear rise and decay envelopes. The individualized SNR levels were defined as the point where participants reached approximately 85% accuracy, ranging from −16dB to 2dB for tone tasks and −12dB to 12dB for consonant tasks.

The following three formal experimental sessions were theta band (4 Hz) tACS, gamma band (40 Hz) tACS, and a sham session (Fig. 2*A-C*). The order of formal sessions was counterbalanced across participants using a balanced Latin square procedure, with a between-session gap of at least five days. Online tACS was applied during the experiment in each session. In sham conditions, 30-second’ short currents were only applied during the ramp-up and ramp-down periods before and after the experiment. Each session consisted of four blocks: lexical tone/consonant identification in either a quiet or noisy environment. Block order was counterbalanced across participants and sessions using a balanced Latin square design. One block included four mini-blocks, with each mini-block randomly presenting all syllables in the continuum. To improve the reliability of slope fits, extra trials near the PSE were included in each mini-block, resulting in a total of 156 trials (39 trials per mini-block). During each trial, participants listened to a syllable and made a speeded perceptual decision via pressing the “<” or “>” keyboard key with their left or right index or middle fingers to judge the lexical tone (tone1 or tone2) or consonant (“d” or “t”) of the heard syllable. Participants’ responses and reaction times were recorded. The responding hand and key assignments were counterbalanced across participants. A maximum 5-second resting period was allowed after sound onset for participants to respond. The next trial began automatically 0.5 seconds (range: 0.4–0.6s) after the previous response or the trial’s end.

Before each formal session, participants completed practice tasks to familiarize themselves with the experimental procedure. The practice task before the first session involved brief tone/consonant identification tasks using a different continuum matrix ([p]–[p^h^] × tone1–tone2 in Mandarin), whereas the practice tasks before the second and third sessions included a short version of the formal experiment. At the end of each electric stimulation, participants rated the intensity of the perceived sensation on a 4-point scale (see Supplementary Materials, Q4), with 1 and 4 indicating no sensations and strong sensations, respectively (No significant differences were found among three tACS conditions: *F*(2, 54) = 1.87, *p* = 0.16; Gamma: mean = 16.26, SD = 3.62; Theta: mean = 15.42, SD = 3.75; Sham: mean = 14.31, SD = 3.61).

### Statistical analysis

#### Fitting psychometric functions

First, we applied binary coding to participants’ responses (i.e., 0 for “d”/“tone1” and 1 for “t”/“tone2”) in the consonant/lexical tone perception task and fitted a four-parameter logistic models using the nonlinear *lsqcurvefit* function in Matlab R2016a, with morphing steps in a continuum as the explanatory variable and responses as the response variable. The slope of the fitted curve at the mid-point served as an index of perceptual sensitivity (Kingdom & Prins, 2016). The formula of this curve is shown below:

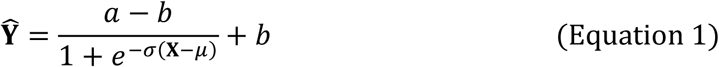

Here, **Ŷ** is the estimated psychometric function (identification curve). **X** is the steps of a continuum. *α* and *b* are the upper and lower asymptotes of the curve. *μ* is the PSE. *σ* is the slope of the curve at the PSE. Slopes of psychometric functions served as the response variable. The lower the slopes, the weaker the participant’s capacity to categorize phonemes. A four-parameter logistic function was employed instead of a simplified model with asymptotes fixed at 0 and 1. This decision was motivated by two main considerations. First, it was unrealistic to assume perfect performance (i.e., 100% accuracy) even when the tone or consonant was highly discriminable, particularly under noise-masking conditions. Second, to mitigate overfitting, the upper and lower asymptotes for each condition were determined by an unconstrained fit to the average data and were subsequently fixed for each block. For each condition, slopes outliers exceeding three standard deviations (SD) from the mean were excluded. However, given that tACS may exert inhibitory effects and potentially induce fitting failure, we adopted a correction procedure for participants with a valid slope in sham but an outlier slope in tACS conditions. Specifically, the outlier slope was replaced with a floor value derived from the corresponding sham blocks (Liang et al., 2023). This floor value was computed as: F = exp(mean(log(D)) −2*SD(log(D))), where D was the slope estimates in sham conditions. We adopted this step to minimize bias introduced by outliers while preserving potential tACS effects. Subsequently, both sham and adjusted tACS slope values were then log-transformed to correct for skewness. To assess the statistical significance of tACS effects for each task (tone/consonant perception under quiet/noise conditions), we conducted a permutation test. Participants’ slope values were shuffled between tACS and sham blocks. For each iteration, the mean difference between conditions was calculated. This procedure was repeated 10^5^ times to generate a null distribution of mean differences. The *p*-value was determined by ranking the observed effect within the permutation distribution, using the following formula: *p* = (ranking + 1) / (10^5^ +1).

### Linear Mixed-Effects Modeling of Reaction Times

We then tested whether tACS significantly altered reaction times (RTs) in each condition by using linear mixed-effects (LME) models. To address the first question and test three associated hypotheses, we established a global model (Model 1) to comprehensively examine the effects of tACS on RTs under different task types and environments (quiet or noisy). The fixed effects in the model included *Stimulus type* (theta vs. gamma tACS), *Task type* (tone vs. consonant perception), and *Environment* (quiet vs. noisy). The *Participant* was included as a random intercept. To focus on testing for tACS effects and to avoid model convergence problems, no random effects were modelled for slope in this study. The global model for RTs is specified as follows:

RTs ∼ Stimulus type * Task type * Environment + (1| Participant) (Model 1)

Given that tone and consonant are two distinct speech cues that may rely on different processing mechanisms (i.e., involving different neural oscillation patterns) (Rufener, Zaehle, et al., 2016; Meyer, 2018), it is worthy examining the effects of tACS separately for the two *Task types* (tone vs. consonant perception). Therefore, two sub-models (Model 2) for tone and consonant tasks were constructed to investigate further the interactions between tACS and the acoustic environment, respectively. Model 2 removed the *Task type* factor from the original model (Model 1). In these models, *Stimulus types* and *Environment* were included as fixed effects, with the *Participant* modeled as a random intercept. The best models for RTs are specified as follows:

RTs ∼ Stimulus type * Environment + (1| Participant) (Model 2 Tone) RTs ∼ Stimulus type * Environment + (1| Participant) (Model 2 Consonant)

All models were implemented using the lme4 package in RStudio 1.4.1103 (R version 4.0.4), and model tests were conducted using the lmeTest package. P-values for the fixed effect (tACS) slopes were corrected for multiple comparisons using the false discovery rate (FDR) method across all tACS conditions within each task.

### Hierarchical Drift-Diffusion Modeling

To gain deeper insight into the cognitive mechanisms through which tACS may influence Mandarin syllable perception, particularly in terms of how tACS modulates decision-related processes beyond surface-level behavioral measures, we further applied a hierarchical drift-diffusion model (HDDM) to fit the pooled trial-level RT distributions across all participants (Wiecki et al., 2013). This model describes perceptual decision-making as an evidence accumulation process modulated by four key parameters: *a* (boundary threshold), *v* (drift rate), *z* (starting point), and *t* (non-decision time). The analyses were conducted using the HDDM package (version 0.9.8) based on Python 2.7.

To test whether tACS significantly affected the parameters *a*, *v*, and *z*, we constructed two linear regression models for each task (tone vs. consonant perception under quiet and noise conditions), assessing the effects of 40 Hz and 4 Hz stimulation separately. *Stimulus type* was dummy-coded with the sham condition serving as baseline. The probability that the parameter posterior distributions deviated from zero (two-tailed test) reflected the confidence (the *p* value) in rejecting the null hypothesis (i.e., no effects of tACS). All *p*-values were then FDR-corrected for multiple comparisons across all tACS conditions. Although non-decision time (*t*) was included in model fitting, it was treated as an intercept-only parameter to avoid overfitting and to prevent overinterpretation of its psychological significance (Weindel et al., 2021).

### Data and code accessibility

Data and code are available on request.

## Results

### Theta tACS impairs consonant sensitivity in quiet

To assess how tACS influenced speech perception, we first analyzed psychometric functions across all experimental conditions (Fig. 3). The slope of each function served as an index of perceptual sensitivity for tone and consonant categorization (Fig. 3). Critically, for consonant perception in the quiet condition (Fig.3*A*), theta tACS significantly lowered the psychometric slope (compared with sham: *p_fdr_* = 0.031, Cohen’s *d* = −0.499), indicating a decrease in perceptual sensitivity; in contrast, gamma tACS did not significantly affect the slope (compared with sham: *p_fdr_* = 0.216, Cohen’s *d* = −0.231), and only marginal significant difference was found between theta and gamma tACS conditions (gamma > theta: *p_fdr_* = 0.083, Cohen’s *d* = 0.36). However, neither theta nor gamma tACS affected slope values for tone perception in quiet (all *p* > 0.05, Fig.3*B*). When noise was added, tACS had no significant effects on psychometric slopes in either tone or consonant perception (all *p* > 0.05, Fig.3*C-D*).

**Figure 3.**
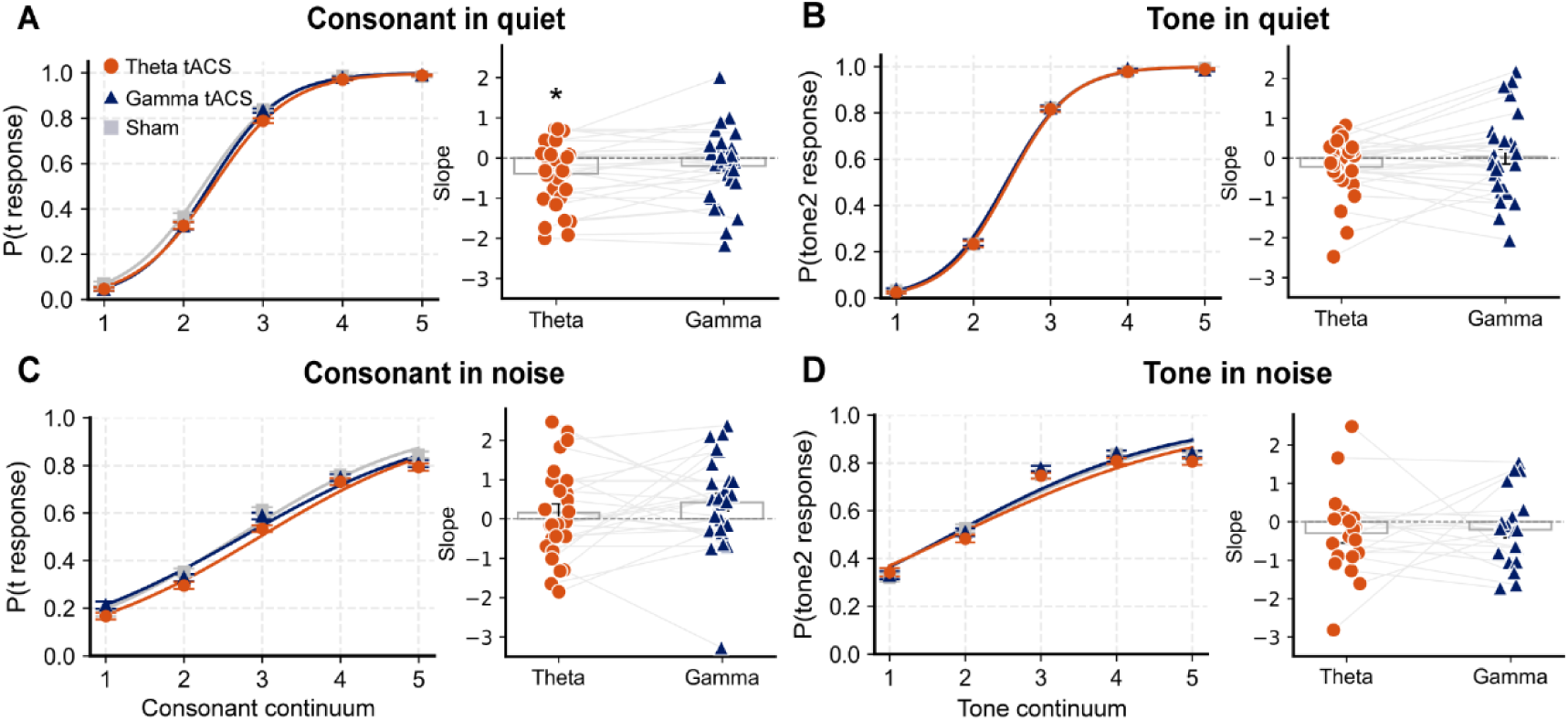
Sigmoid psychometric functions and corresponding slope estimates at the points of subjective equity (PSE, step3) across five-step continua in different tACS and listening conditions. ***A-D***, Each panel contains two plots: the left plot shows the psychometric function (proportion of categorical responses across continuum steps), and the right plot shows the PSE slope of the function for each stimulation condition. ***A***-***B,*** In quiet conditions, the proportions of “t” responses (***A***, consonant continuum) or “tone 2” responses (***B***, tone continuum) were plotted along continuum steps. ***C***-***D***, The same measures were shown under background noise. Lines in the left sub-plots of each panel indicate mean responses across participants for each of the three stimulation conditions (theta tACS: orange; gamma tACS: blue; sham: gray), with error bars representing standard error. All psychometric functions reflect increased “t” or “tone 2” responses from acoustic step 1 to 5. Noted that, in ***A***, theta tACS elicited a shallower slope compared to gamma tACS and sham, suggesting a reduced sharpness of the category boundary under theta stimulation. Compared with sham, only theta tACS lowered the psychometric slope in the consonant task in quiet (\**p_fdr_* = 0.031, two-tailed paired t-test). No observable differences in slope were observed between stimulation conditions in tone perception (***B***, ***D***) or in the consonant in noise condition (***C***).

In the right subplots of each panel, each dot represents one participant; gray lines connect paired data points across stimulation conditions. Bars indicate group means.

We next examined whether tACS altered response bias by analyzing the PSE in consonant categorization (see Figure S1). In the quiet conditions, both gamma and theta tACS induced marginal shifts in PSE relative to sham (gamma: *p_fdr_* = 0.081, Cohen’s *d* = 0.298; theta: *p_fdr_* = 0.081, Cohen’s *d* = 0.325), hinting at a potential, albeit weak, modulation of categorical boundaries. A similar trend was observed for theta tACS in noise (compared with sham: *p_fdr_* = 0.083, Cohen’s *d* = 0.430), suggesting that theta stimulation might subtly bias perceptual thresholds even in challenging listening environments. However, no significant PSE effects were found for tone perception in either quiet or noise conditions (all *p* > 0.05).

In summary, the psychometric effects of tACS were mainly observed in consonant perception under quiet conditions: theta tACS significantly reduced perceptual sensitivity (slope decrease, Fig. 3*A*), suggesting impaired encoding of phonemic contrasts. Marginal shifts in PSE were also observed for both theta and gamma tACS, but only in consonant categorization. These findings indicate that theta-band neural oscillations are causally involved in phoneme boundary perception in Mandarin. Such effects were primarily observed in consonant categorization under quiet listening conditions.

### Theta tACS prolongs RTs across tasks and environments but gamma only delays tone judgment in quiet

We then proceeded to comprehensively characterize the temporal dynamics of tACS effects using LME models (Table 1). This approach allowed us to account for both fixed effects of experimental conditions and random variations across participants while examining RT differences. The results are reported in an analysis of variance (ANOVA) format.

**Table 1.** Modulations of tACS effects on RTs in different conditions (seconds, *Mean* ± *SD*).

| Stimulus type | In quiet |  | In noise |  |
| --- | --- | --- | --- | --- |
|  | Consonant | Tone | Consonant | Tone |
| Theta tACS | 0.77±0.24 | 0.81±0.23 | 0.88±0.27 | 0.96±0.33 |
| Gamma tACS | 0.76±0.22 | 0.80±0.23 | 0.86±0.27 | 0.94±0.33 |
| Sham | 0.75±0.24 | 0.79±0.24 | 0.86±0.27 | 0.95±0.36 |

In Model 1, a significant main effect of *Stimulus type* was observed (χ²(2) = 54.480, *p* < 0.001). Post-hoc tests demonstrated that theta tACS significantly prolonged RTs relative to both sham (*β* = –0.017, *SE* = 0.002, *t* = –6.998, *p* < 0.001) and gamma tACS (*β* = –0.013, *SE* = 0.002, *t* = –5.522, *p* < 0.001), whereas gamma tACS did not differ significantly from sham (*β* = –0.003, *SE* = 0.002, *t* = –1.479, *p* = 0.139). A significant main effect of *Task type* emerged (χ²(1) = 893.890, *p* < 0.001), with longer RTs observed for tone compared to consonant categorization (*β* = –0.059, *SE* = 0.002, *t* = – 29.915, *p* < 0.001). Additionally, RTs were strongly modulated by *Environment* (χ²(1) = 4389.810, *p* < 0.001), with longer RTs observed in noise versus quiet conditions (*β* = –0.130, *SE* = 0.002, *t* = –66.262, *p* < 0.001).

A significant interaction was found between *Stimulus type* and *Environment* (χ*²*(2) = 6.410, *p* = 0.041), indicating that tACS effects were modulated by task difficulty. Simple effects analysis revealed that, in quiet environment, RTs under both theta tACS (*β* = –0.021, *SE* = 0.003, *t* = –6.230, *p* < 0.001) and gamma tACS (*β* = –0.009, *SE* = 0.003, *t* = –2.772, *p* = 0.006) were significantly longer compared to sham, with theta tACS exerting an even stronger effect than gamma (*β* = –0.012, *SE* = 0.003, *t* = –3.458, *p* < 0.001). In noisy environment, theta tACS kept lengthening RTs relative to sham (*β* = –0.013, *SE* = 0.003, *t* = –3.669, *p* < 0.001) and gamma tACS (*β* = –0.015, *SE* = 0.003, *t* = –4.351, *p* < 0.001), whereas no significant difference was observed between gamma tACS and sham (*β* = 0.002, *SE* = 0.003, *t* = 0.679, *p* = 0.497). This pattern suggests that theta tACS consistently impeded response speed, regardless of environmental noise, while gamma tACS showed weaker and condition-specific effects, which are constrained to the unchallenging listening environment.

To examine whether tACS effects were modulated differentially by *Task types*, we analyzed consonant and tone perception tasks separately using the same LME framework (RTs ∼ Stimulus type * Environment + (1| Participant)).

In Model 2 Consonant (Fig. 4*A,C*), significant main effects of *Stimulus type* (*χ²*(2) = 43.641, *p* < 0.001) and *Environment* (*χ²*(1) = 1923.536, *p* < 0.001) were observed. Post-hoc tests showed that theta tACS again slowed RTs versus sham (*β* = –0.019, *SE* = 0.003, *t* = –6.278, *p* < 0.001) and gamma tACS (*β* = –0.015, *SE* = 0.003, *t* = –4.915, *p* < 0.001), while gamma tACS and sham did not differ (*β* = –0.004, *SE* = 0.003, *t* = – 1.366, *p* = 0.172). The interaction between *Stimulus type* and *Environment* was nonsignificant (*χ²*(2) = 1.015, *p* = 0.602).

**Figure 4.**
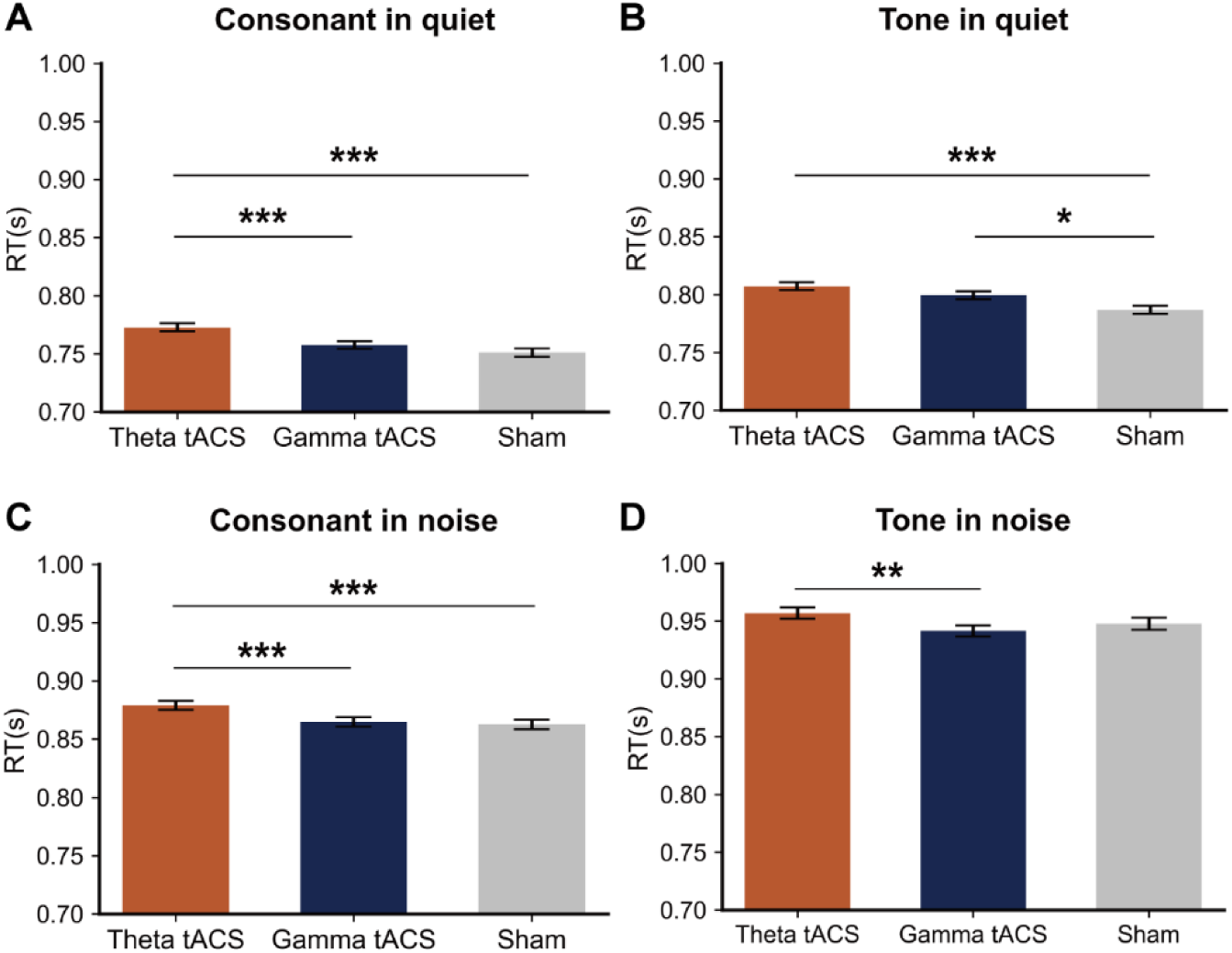
Reaction times (RTs) across tACS conditions in consonant and tone perception tasks in quiet and noise. ***A***, In the consonant tasks in quiet, theta tACS resulted in significantly longer RTs compared to both gamma tACS and sham. ***B***, In the tone tasks in quiet, both stimulation conditions led to longer RTs than sham. ***C***, In the consonant tasks in noise, RTs were significantly prolonged under theta tACS compared to both gamma tACS and sham. ***D***, In the tone tasks in noise, only theta tACS led to significantly longer RTs than gamma tACS. \*\**p_fdr_* < 0.01. \*\*\**p_fdr_* < 0.001. Bars indicate group means; error bars indicate standard error.

In Model 2 Tone (Fig. 4*B,D*), analysis of RTs revealed a significant main effect of *Stimulus type* (*χ²*(2) = 17.933, *p* < 0.001). Post-hoc tests indicated that theta tACS prolonged RTs relative to sham (*β* = –0.015, *SE* = 0.004, *t* = –4.007, *p* < 0.001) and gamma tACS (*β* = –0.012, *SE* = 0.004, *t* = –3.186, *p* = 0.002), while gamma tACS effects did not differ from sham (*β* = –0.003, *SE* = 0.004, *t* = –0.822, *p* = 0.411). A significant main effect of *Environment* was also observed (*χ²*(1) = 2582.679, *p* < 0.001), with RTs being significantly longer in noise conditions compared to quiet conditions (*β* = –0.152, *SE* = 0.003, *t* = –50.820, *p* < 0.001).

The interaction between *Stimulus type* and *Environment* was significant (*χ²*(2) = 6.664, *p* = 0.036). Simple effects analyses in Figure 4*B* showed that, in quiet, both theta (*β* = –0.020, *SE* = 0.005, *t* = –3.950, *p* < 0.001) and gamma tACS (*β* = –0.012, *SE* = 0.005, *t* = –2.392, *p* = 0.025) significantly increased RTs compared to sham, with no significant difference between theta and gamma tACS (*β* = –0.008, *SE* = 0.005, *t* = – 1.558, *p* = 0.119). In noise (Fig. 4*D*), neither theta nor gamma tACS significantly differed from sham (all *p* > 0.05). However, RTs under theta tACS were significantly longer than those under gamma tACS (*β* = –0.015, *SE* = 0.005, *t* = –2.946, *p* < 0.01). Notably, the effect of gamma tACS was only present in the quiet condition.

In sum, these results from both models indicated that theta tACS reliably resulted in longer RTs compared to sham across tasks and environments, suggesting a broad impact on processing efficiency. By contrast, gamma tACS showed more selective effects, increasing RTs mainly during tone perception in quiet. This extends our findings on the psychometric slope by evidencing a role of theta oscillations in temporal integration mechanisms and of gamma oscillations in tone-related auditory processing during Mandarin speech perception.

### Theta and gamma tACS modulate the perceptual decision-making in Mandarin syllables

To dissect what distinct cognitive subprocesses of speech perceptual decision making are supported by theta and gamma neural oscillations, we applied HDDM to account for both responses and reaction RTs to decompose decision making into three interpretable parameters: boundary threshold (*a*) reflecting threshold for evidence accumulation, drift rate (*v*) indexing evidence accumulation rate, and starting point (*z*) capturing initial response bias (Fig. 5). TACS effects on the three parameters were quantified by linear regression (tACS vs. Sham).

**Figure 5.**
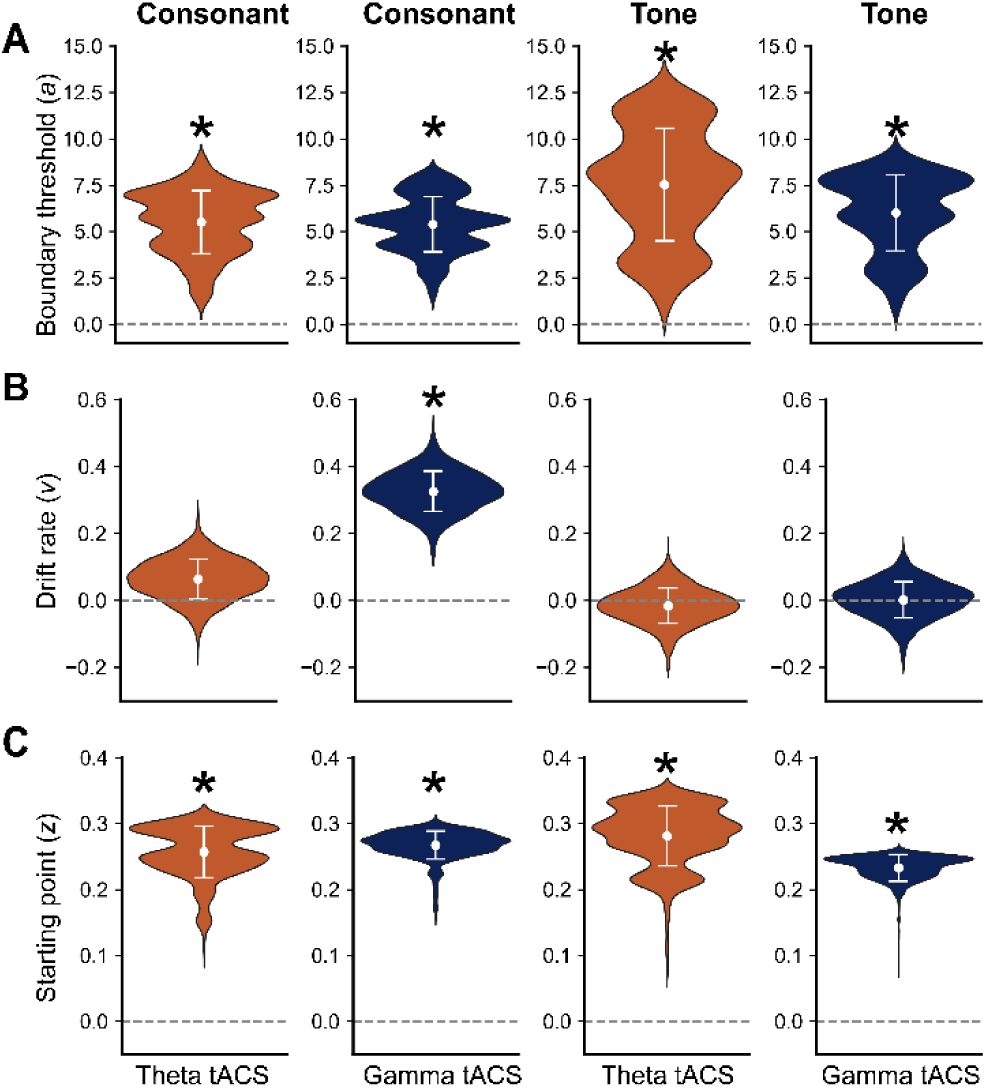
TACS effects on HDDM parameters across tasks. Each row displays violin plots of one parameter estimated from hierarchical Bayesian modeling: boundary threshold (*a*, ***A***), drift rate (*v*, ***B***), and starting point (*z*, ***C***), treating Sham as baseline. Within each sub-plot, data were plotted separately for consonant (left) and tone (right) perception tasks under theta (orange) and gamma (blue) tACS. Error bars represent SEM. \**p_fdr_* < 0.05 (two-tailed). ***A***, Boundary threshold (*a*), indicating the amount of evidence required to trigger a decision, was significantly increased in all conditions. Larger values reflect more cautious decision-making. ***B***, Drift rate (*v*), reflecting the efficiency of evidence accumulation, showed a significant rise only in the consonant task under gamma tACS. Larger values indicate more efficient stimulus-driven processing. ***C***, Starting point (*z*), representing prior bias toward a particular response, significantly deviated from zero in all conditions. Compared with Sham, positive values indicate a prior bias toward choosing /d/ or tone 1; negative values indicate a bias toward /t/ or tone 2.

For consonant perception, gamma tACS significantly elevated the boundary threshold (*a*), indicating that participants required more evidence to reach a decision. It also enhanced the drift rate (*v*), reflecting a faster accumulation of sensory information. Additionally, gamma stimulation shifted the starting point (*z*) toward the “d” response, suggesting a stronger initial bias toward that choice (all *p_fdr_* < 0.001, two-tailed test). Theta tACS similarly raised boundary threshold (*a*) and altered starting point (*z*) (all *p_fdr_* < 0.001, two-tailed test), but did not significantly alter the drift rate (*v*) (*p_fdr_* = 0.144). In the tone perception task, both gamma and theta tACS significantly increased the parameter *a*, with all posterior distributions deviating from zero (all *p_fdr_* < 0.001, two-tailed test). This indicated that participants required more evidence to make a decision after gamma or theta oscillations were perturbed. TACS also significantly perturbed the parameter *z*, suggesting a response bias shift toward tone 1 choice. However, neither gamma (*p_fdr_* = 0.463) nor theta tACS (*p_fdr_* = 0.376) significantly affected the drift rate (*v*), suggesting that the speed of evidence accumulation remained unchanged.

To summarize, theta and gamma tACS consistently increased the boundary threshold (*a*) and biased the starting point (*z*) upward across tasks, suggesting increased caution and altered decision biases. Notably, only gamma tACS led to a significant increase in drift rate (*v*) during consonant perception, indicating increased evidence accumulation speed unique to this condition.

## Discussion

In the current study, we applied theta and gamma tACS to the temporal cortices of Mandarin-speaking participants during different Mandarin phoneme identification tasks and examined their impact on performance. We aimed at probing whether and how neural oscillations in these two frequency bands causally support Mandarin syllable perception, and how the mechanisms are modulated by the distinctive features (i.e., consonant vs. lexical tone) and task difficulty (quiet vs. noisy environment).

### Syllable-level representations in Mandarin phoneme perception supported by theta oscillations

The present study provides causal evidence that theta oscillations play a key role in consonant and lexical tone perception in Mandarin, broadly affecting processing efficiency (Fig. 4). In contrast, gamma oscillations exert more selective influences, particularly in lexical tone perception in quiet conditions. These results largely supported Hypothesis 3, which states that phonemes in Mandarin are identified as holistic syllabic units that encapsulate both suprasegmental and segmental cues, subserved by theta oscillations (Fig. 1*C*). Notably, this effect remained evident for consonant perception even under noisy conditions (Fig. 4*C*).

Theta oscillations have been found to track speech rhythm and integrate slowly varying suprasegmental features in speech (Meyer, 2018; Teoh et al., 2019). In Mandarin, where the syllable serves as a fundamental perceptual unit, the role of theta rhythms in holistic syllable processing appears especially critical (He et al., 2023; Ni et al., 2023). Our results suggest that the functions of theta oscillations are not limited to suprasegmental tone integration but extend broadly to segmental phonemes (e.g., consonants). This finding contradicts similar tACS studies on English syllable identification, which have found that gamma tACS selectively modulates the perception of segmental phonemes (Marchesotti et al., 2020; Rufener, Oechslin, et al., 2016; Rufener, Zaehle, et al., 2016), corresponding to their modulation rates at around 40Hz. However, in our current study, theta, but not gamma, tACS impaired the perception of consonants embedded in Mandarin syllables. One explanation is that Mandarin evolved from Old Chinese, which was monosyllabic, and retains strong syllable-word or syllable-morpheme mappings (Feng, 2015). Moreover, Mandarin contains a large corpus of homophones(Huang & Liao, 2017; Li et al., 2024), reflecting its lower diversity in phoneme combinations, or in other words, higher inter-phoneme predictability within a syllable, than English. The “monosyllabic” feature of modern Mandarin can lead to syllables being processed by the cognitive system as a whole unit, rather than as a combination of discrete phonemes.

Moreover, evidence from event-related potential (ERP) study (Zhao et al., 2011; Zou et al., 2020) and phonetic research (Kang & Xu, 2024; Liu & Xu, 2023) has shown a high degree of synchronization between tone and vowel processing, supporting the holistic view of syllable perception. Thus, the effects of theta tACS on Mandarin consonant identification can reflect that although participants needed to distinguish subtle differences in segmental phonemes (e.g., VOT), the phonemes themselves might be represented as syllables and are therefore more dependent on slow theta oscillations. An alternative explanation is that the experiment itself used meaningful words that could trigger holistic semantic processes. However, another study using the same experimental task and materials selectively modulated the perception of lexical tone and consonants with transcranial magnetic stimulation, which suggests that this task can effectively elicit phoneme-level representations (Liang et al., 2023). Additionally, we used natural Mandarin syllables to maintain high ecological validity.

Importantly, although theta tACS effects on psychophysical sensitivity were attenuated under noisy conditions or even absent (Fig. 3*A, C*), its inhibitory influence on RTs in both consonant and tone tasks persisted (Fig. 4). This pattern suggests that noise, as a perceptual interference, may alter the endogenous expression of theta oscillations or modulate their interaction with externally applied tACS to perceptual sensitivity, possibly through increased perceptual load (Yellamsetty & Bidelman, 2018). Nevertheless, syllable-based auditory representations in Mandarin phonemes remain essential even in challenging listening tasks. However, theta tACS on the auditory cortices did not alter the perceptual sensitivity under noise conditions, potentially indicating lower auditory processing but stronger motor compensation supported by oscillations at higher frequency bands (e.g., alpha or beta (Jenson et al., 2014; Saltuklaroglu et al., 2018)) instead.

### Theta and gamma oscillations support the perceptual decision in Mandarin syllables

Results from the HDDM revealed that theta tACS systematically increased the decision boundary (*a*, Fig. 5*A*) and modified the starting point (*z*, Fig. 5*C*). Besides hampering the auditory sampling and impairing speech perception (Meyer, 2018), indicated by the increased amount of perceptual evidence required for decision (*a*), theta tACS also affected the response bias (*z*) that determines the decision relying more on memory instead of the upcoming speech input. Meanwhile, since we applied random inter-trial intervals, it is unlikely that this response bias was caused by tACS-induced phase resetting of neural oscillations (Riecke et al., 2015), which would lead to a systematic change in the sampling of VOT and pitch contour. Therefore, the modulation of response bias by theta tACS may reflect the role of theta oscillations in supporting memory functions or general motor decision-making processes; however, future studies are needed to draw a solid conclusion.

More importantly, gamma tACS also exerted similar effects on the *a* and *z* as theta tACS did, while no significant gamma tACS effects on the psychometric slopes and reaction times (except the RTs in lexical tone identification in quiet) were found. This indicates that gamma oscillations are also recruited in Mandarin syllable processing, but the recruitment tends to be compensatory rather than necessary. Indeed, gamma oscillations are critically involved in processing local, rapidly changing speech features (Fukuda et al., 2010; Lizarazu et al., 2019). Specifically, under quiet conditions, gamma activity may retain its advantage in processing fine-grained acoustic features, particularly pitch cues (Yellamsetty & Bidelman, 2018), which are less susceptible to noise. This corresponds to our finding that gamma tACS prolonged lexical tone identification but only in quiet. Indeed, one recent EEG study shows that the perception of lexical tone depends on temporal fine structure, supporting our results (Ni et al., 2023). On the other hand, gamma tACS significantly enhanced the drift rate (*v*) for consonant processing. Noted that, gamma tACS did not modulate the psychometric slopes or reaction times in consonant identification, regardless of environment. Previous studies showed that changes in drift rate indicate that top-down predictive processes enhance the rate of evidence accumulation. Thus, it is plausible that gamma oscillations were in fact actively supporting consonant perception, and hence were impaired by the gamma tACS (also indicated by the altered *a* and *v*). However, compensation from the higher levels of representations (e.g., syllable level) may counteract the inhibited, gamma-based, temporally fine phoneme encoding.

In brief, gamma tACS modulated the HDDM parameters of perceptual decisions like theta tACS, but only affected the performance of tone perception in quiet. This indicates that gamma oscillations are also involved in Mandarin phoneme perception, supporting the encoding of fast-changing features with fine-grained spectrotemporal patterns. However, their role is modulatory rather than causal, as the theta-supported holistic syllable representation can easily complement the degraded fine representations induced by impaired gamma activity.

### Limitations

Several limitations could affect the interpretations derived from the current results. First, this study did not directly measure neural oscillations, especially the modulatory effects of tACS. However, previous studies have shown that in theta and gamma tACS can modulate the corresponding neural oscillations (Ghiani et al., 2021; Meyer, 2018), and a link between neural perturbations and behavioral changes has also been provided (Riecke et al., 2015; Rufener, Zaehle, et al., 2016). Second, in this study, we only recruited Mandarin-speaking participants to complete Mandarin identification tasks. Future studies are recommended to directly compare the perception of tonal and non-tonal languages (e.g., Mandarin vs. English) within and across groups of participants from different language backgrounds.

### Conclusions

Collectively, the theta tACS effects observed in this study support our third hypothesis, “syllable-level holistic processing” which posits that the auditory encodings of Mandarin phonemes (including suprasegmental lexical tones and segmental consonants) are embedded in holistic syllable representations, supported by theta oscillations that facilitate the temporal sampling of syllabic cues. As tACS has been postulated to reveal causal effects by directly modulating neural activity (Herrmann et al., 2016), our study further implicates the causal role of theta oscillations in the perception and identification of Mandarin consonants and lexical tones, regardless of task difficulty. Furthermore, our findings indicate that gamma oscillations are also engaged, but with a modulatory role, in resolving fine-grained spectrotemporal acoustic patterns, which can be complemented by the theta-based holistic syllable representations. Therefore, we propose that theta oscillations dominate Mandarin consonant and lexical tone perception through holistic syllable processing, but also with gamma compensation.

## Supplementary Materials

**Questionnaire 1. Transcranial Alternating Current Stimulation (tACS)**

### Experiment Volunteer Screening Questionnaire

[This questionnaire is intended to screen potential participants for a behavioral experiment. Only currently enrolled university students are eligible to complete this form.]

If you agree to participate in this experiment, please answer the following questions. The information you provide will be used solely for screening purposes and will be kept strictly confidential.

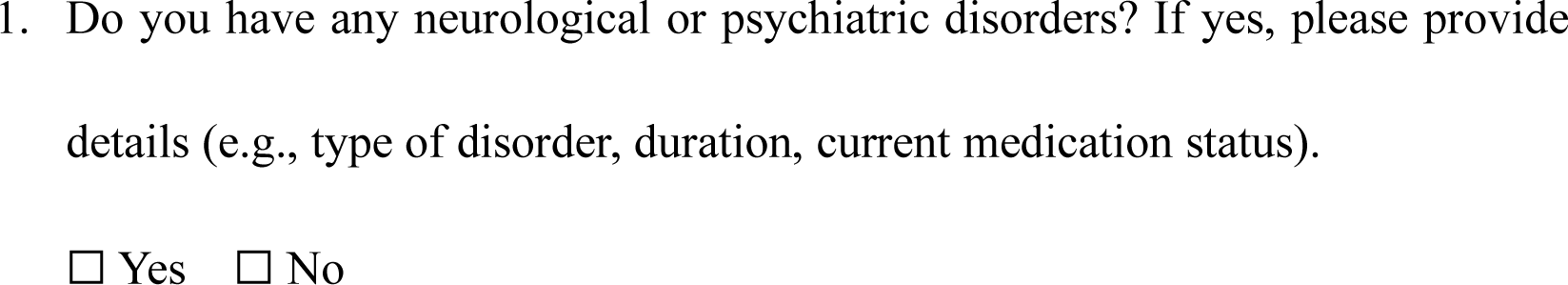
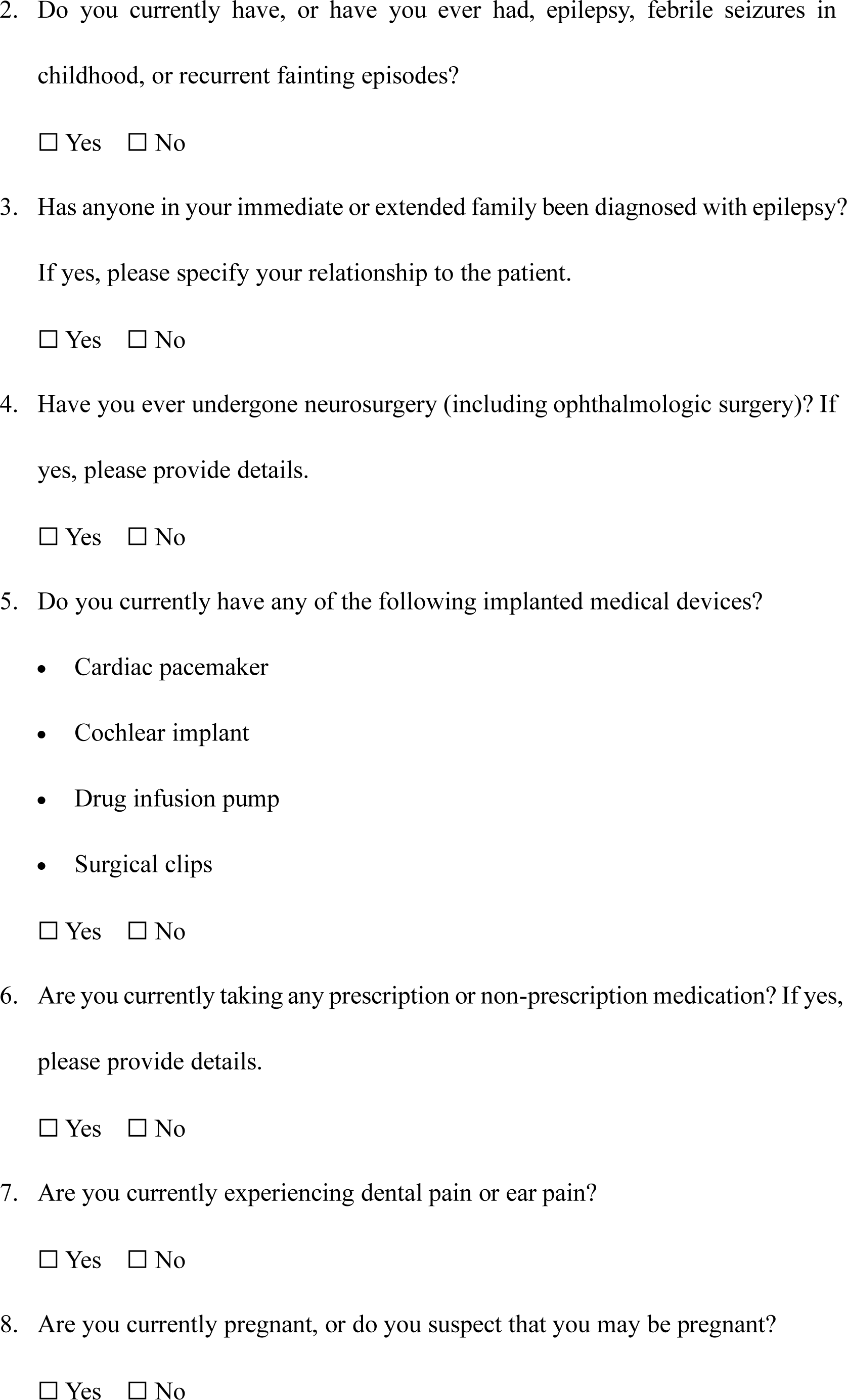
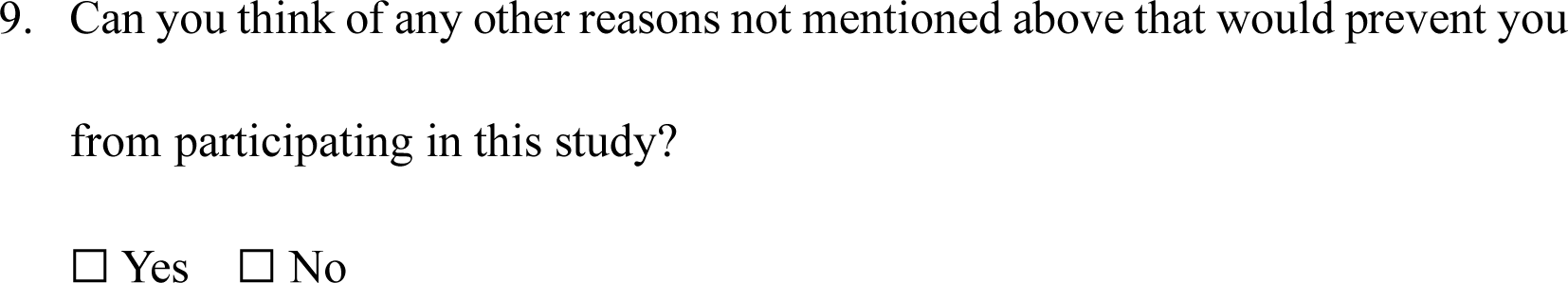

**Questionnaire 2. Musical Training Experience Questionnaire**

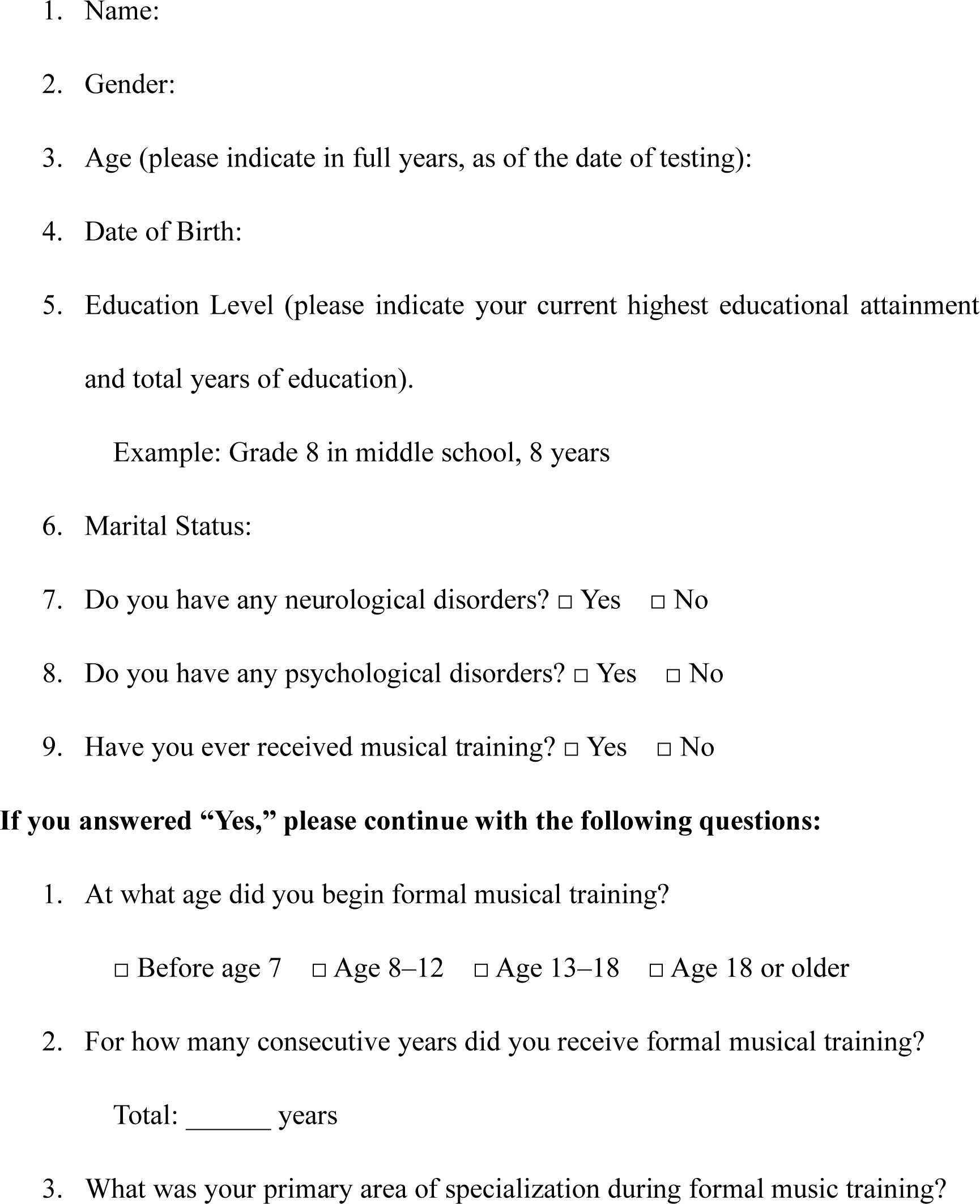
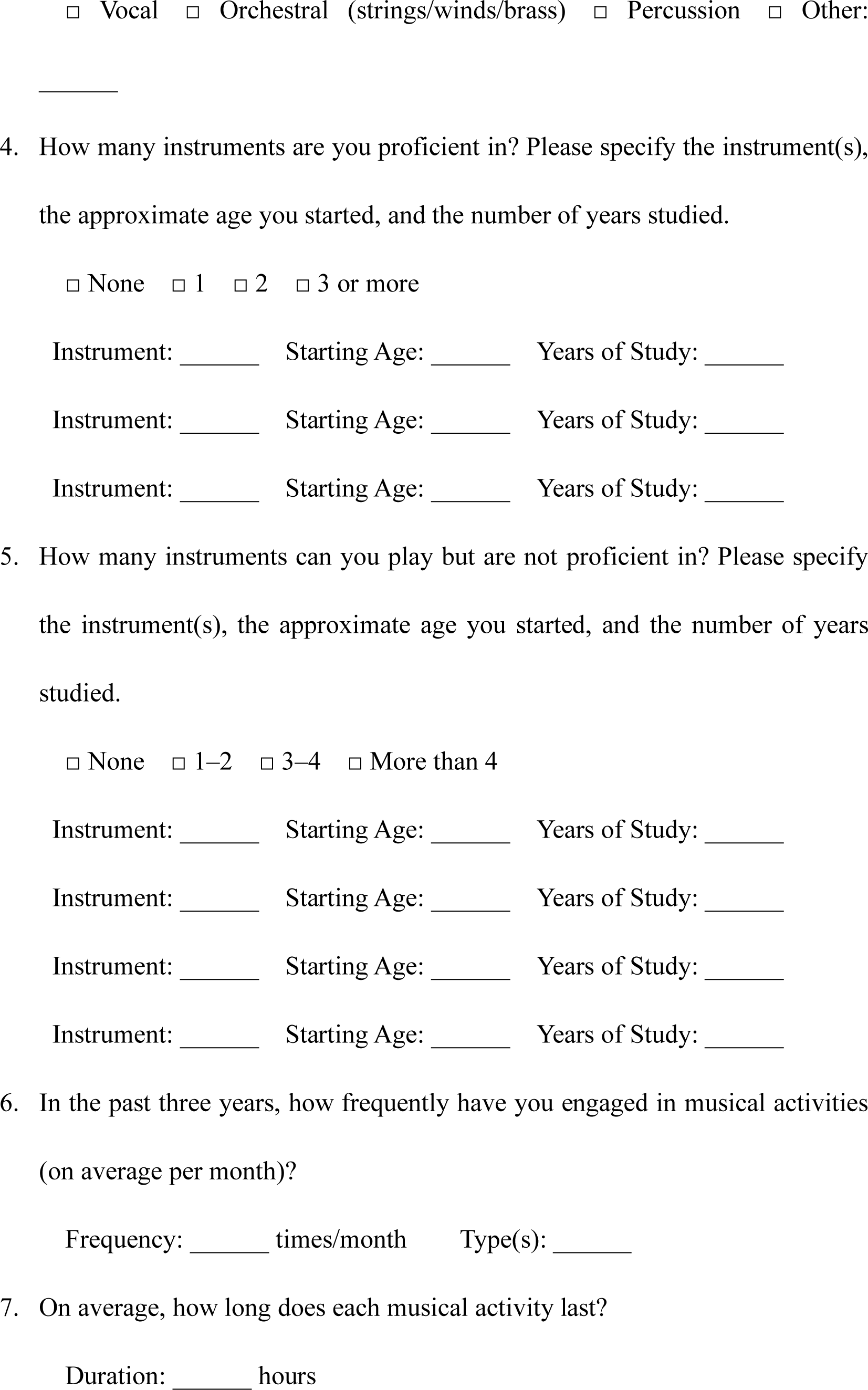

**Questionnaire 3. Edinburgh Handedness Inventory^1^**

For each of the following tasks, please select the hand you **prefer** to use.

- If your preference is very strong, such that you would never use the other hand unless forced, please select **Left hand** or **Right hand**.
- If there is no clear preference and you use **both hands equally**, please select

Both hands.

- Some activities naturally require both hands. In such cases, please judge only the hand preference for the object indicated in parentheses.

| Task / Object | Left hand | Both hands | Right hand |
| --- | --- | --- | --- |
| 1. Writing | ☐ | ☐ | ☐ |
| 2. Drawing | ☐ | ☐ | ☐ |
| 3. Throwing | ☐ | ☐ | ☐ |
| 4. Scissors | ☐ | ☐ | ☐ |
| 5. Toothbrush | ☐ | ☐ | ☐ |
| 6. Knife (without fork) | ☐ | ☐ | ☐ |
| 7. Spoon | ☐ | ☐ | ☐ |
| 8. Broom (upper hand) | ☐ | ☐ | ☐ |
| 9. Match (holding the match) | ☐ | ☐ | ☐ |
| 10. Opening a box (lid) | ☐ | ☐ | ☐ |

Scoring:

- Left hand = –2 points
- Both hands = 0 points
- Right hand = +2 points

Calculation:

- **LH =** total left-hand points
- **RH =** total right-hand points
- **CT =** LH + RH (cumulative total)
- **D =** RH – LH (difference score)
- **R =** (D / CT) × 100 (laterality quotient)

Interpretation:

1. Left-handed: R < –40
2. Ambidextrous: –40 ≤ R ≤ +40

• Right-handed: R > +40

*Reference:*

Oldfield, R. C. (1971). *The assessment and analysis of handedness: The Edinburgh Inventory*. Neuropsychologia, 9, 97–113.

**Questionnaire 4. Post-Stimulation Sensation Questionnaire**

1. **Participant ID** (Format: Serial number + Initials in uppercase. Example: N01XXX. The serial number will be provided by the experimenter.)

2. **Session Number**:

☐ Pilot Session

☐ Formal Session 1

☐ Formal Session 2

☐ Formal Session 3

3. During the experiment, did you experience any of the following symptoms?

Please rate only symptoms that are related to electrical stimulation (e.g., if a headache was due to fatigue and not stimulation, do not assign a high score).

|  | Scale: 1 = None 2 = Mild 3 = Moderate 4 = Severe |
| --- | --- |
| Headache |  |
| Neck pain |  |
| Scalp pain |  |
| Scalp injury |  |
| Skin redness / heating sensation |  |

4. During the experiment, did you experience any of the following symptoms?

Please rate only symptoms that are related to electrical stimulation (e.g., if sleepiness was due to fatigue and not stimulation, do not assign a high score).

|  | Scale: 1 = None 2 = Mild 3 = Moderate 4 = Severe |
| --- | --- |
| Sleepiness |  |
| Difficulty concentrating |  |
| Sudden mood changes |  |
| Dizziness |  |

5. During the experiment, did you experience any of the following visual symptoms? Please rate only symptoms that are related to electrical stimulation (e.g., if blurred vision was due to fatigue and not stimulation, do not assign a high score).

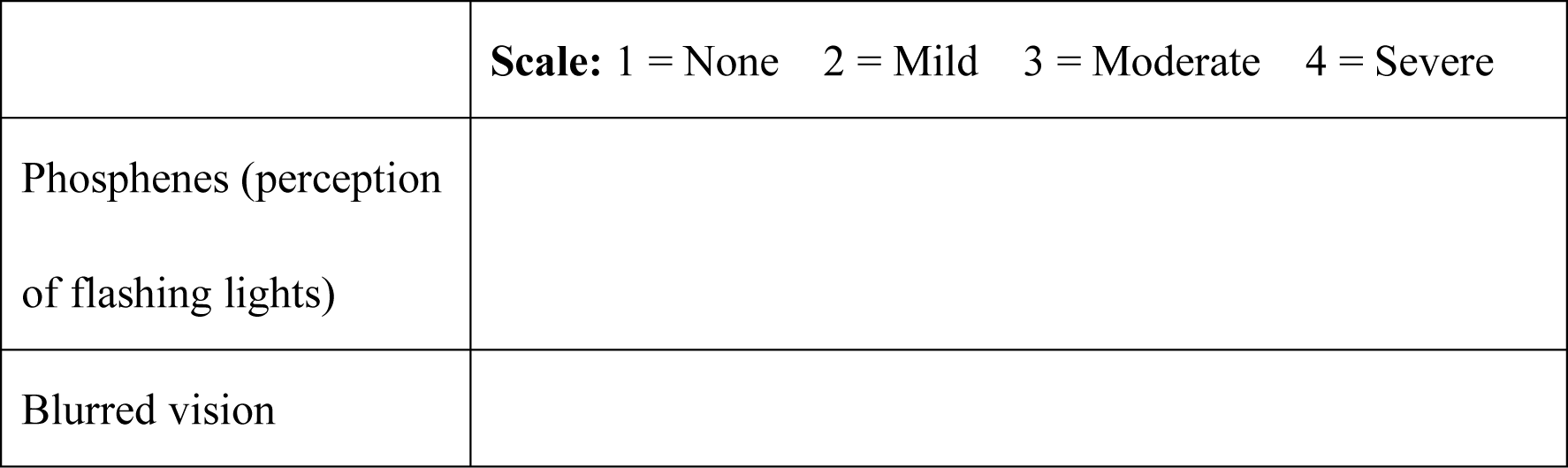

6. During the experiment, did you experience the following symptom?

|  | **Scale:** 1 = None 2 = Mild 3 = Moderate 4 = Severe |
| --- | --- |
| Sensation of increased weight or pressure around the electrode sites or on the head |  |

7. Other (please specify):_____

**Supplementary Results**

**Simple effects analysis following the significant *Task type* × *Environment* interaction in LME model 1**

An additional significant interaction was observed between *Task type* and *Environment* (*χ²*(1) = 118.90, *p* < 0.001). Simple effects analysis indicated that RTs in the tone perception task were significantly longer than those in the consonant perception task, both in quiet (*β* = –0.0374, *SE* = 0.00278, *t* = –13.452, *p* < 0.001) and in noise (*β* = –0.0804, *SE* = 0.00279, *t* = –28.843, *p* < 0.001). All other interactions were not statistically significant (all *p* > 0.05).

## Supplementary Figure

**S1.**
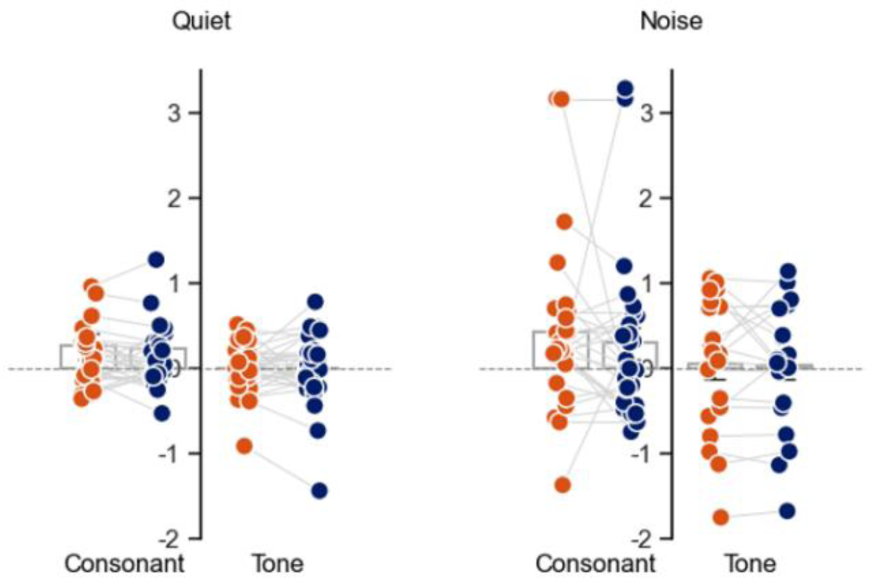
Points of subjective equity. (PSEs) of the psychometric functions for consonant and tone identification tasks under theta and gamma tACS in quiet and noise conditions. Bars indicate group means; error bars indicate SEM. No significant PSE effects were found (*p*s > 0.05).

